# Multiplexed transcriptome discovery of RNA binding protein binding sites by antibody-barcode eCLIP

**DOI:** 10.1101/2022.06.08.495357

**Authors:** Daniel A. Lorenz, Kylie A. Shen, Hsuan-Lin Her, Katie Rothamel, Kasey R. Hutt, Allan C. Nojadera, Stephanie C. Bruns, Sergei A. Manakov, Karen B. Chapman, Gene W. Yeo

**Affiliations:** Eclipse Bioinnovations, San Diego, CA; Department of Cellular and Molecular Medicine, University of California San Diego, La Jolla, CA; Institute for Genomic Medicine, University of California San Diego, La Jolla, CA; Stem Cell Program, University of California San Diego, La Jolla, CA

## Abstract

UV cross-linking and immunoprecipitation (CLIP) methodologies enable the identification of RNA binding sites of RNA-binding proteins (RBPs). Despite improvements in the library preparation of RNA fragments, the current enhanced CLIP (eCLIP) protocol requires 4 days of hands-on time and lacks the ability to process many RBPs in parallel. We present a new method termed antibody-barcode eCLIP (ABC) that utilizes DNA-barcoded antibodies and proximity ligation of the DNA oligonucleotides to RBP-protected RNA fragments to interrogate multiple RBPs simultaneously. We observe performance comparable to eCLIP with the advantage of a reduced hands-on time of 2 days and dramatically increased scaling while minimizing sample-to-sample variation and maintaining the same material requirement of a single eCLIP experiment.

## Main Text

RNA-binding proteins (RBPs) are critical regulators of gene expression, controlling the rate, location, and timing of RNA maturation^1–4^. As such, dysregulation of RBP function is associated with diverse genetic and somatic disorders, such as neurodegeneration and cancer^5,6^. To uncover the molecular mechanisms by which RBPs affect RNA processing, technologies such as RNA immunoprecipitation (RIP) and CLIP coupled with high-throughput sequencing enable the transcriptome-wide identification of RNA binding sites^7–10^. Recent improvements to CLIP library preparation have led to more reproducible and robust CLIP datasets with a higher recovery rate of successful libraries^11–14^. For instance, enhanced CLIP (eCLIP) improvements enabled the generation of 223 eCLIP datasets profiling targets for 150 RBPs in K562 and HepG2 cell lines via a standardized protocol^15^. As part of the ENCODE III phase, these target maps expanded the catalog of functional RNA regulatory elements encoded in the human genome and revealed unexpected principles of RNA processing^14–16^. However, the number of protein-coding genes with experimental or computational evidence for RNA binding properties have continued to increase, accounting for at least ~15% of the human genome^17–20^, and our ENCODE pilot still represents less than 10% of annotated RBPs.

We opine that reducing the technical complexity of the eCLIP protocol is pivotal to accelerate our progress toward an exhaustive characterization of RBPs. While eCLIP has improved library generation efficiency, two major limitations to scaling remain. First, all current CLIP-based methods feature SDS-PAGE and nitrocellulose membrane transfer step to size-select for the immunoprecipitated protein-RNA complex^21–23^. The nitrocellulose membrane captures and separates RNA fragments cross-linked to the protein of interest from free, unbound RNA. However, this manual excision of estimated protein-RNA bands is tedious, requires an additional 1.5-2 days, and is vulnerable to a large degree of user-to-user variation. Second, each individual RBP requires a separate immunoprecipitation (IP) step, which places a burden on the quantity of input material required for studying many RBPs.

Here, we develop antibody-barcode eCLIP (ABC) based on modifications to the eCLIP protocol. Our optimizations address both of eCLIP’s constraints through the incorporation of DNA-barcoded antibodies that allow on-bead proximity-based ligations to replace the SDS-PAGE and membrane transfer steps. DNA-barcoded antibodies have been utilized to adapt protein detection into sequenceable readouts^24^. Here, we utilize the barcodes to distinguish the identity of different RBPs within the same sample, mitigating the input quantity limitation. These modifications shorten the hands-on time by ~1.5 to 2 days (**Fig.1a**). We evaluated ABC using two well-characterized RBPs, RNA Binding Fox-1 Homolog 2 (RBFOX2), which recognizes GCAUG motifs, and the Stem-Loop Binding Protein (SLBP), which interacts specifically with histone mRNAs. Replicate ABC experiments for both RBPs were performed in HEK293T and K562 cells, respectively, and reads were mapped and processed as previously described^13,15^. We compared the library complexity as a surrogate measure of library efficiency by enumerating the number of ‘usable’ reads, defined as reads that map uniquely to the genome and remain after discarding PCR duplicates, as a function of sequencing depth and observed similar library complexity for RBFOX2 eCLIP and ABC (Supplemental Fig.1). Examination of individual binding sites revealed comparable read density between ABC and eCLIP at RBFOX2 (e.g., intronic region of NDEL1) and SLBP (e.g., 3’UTR of H1-2) binding sites (**Fig.1b**)^14^.

**Figure. 1:**
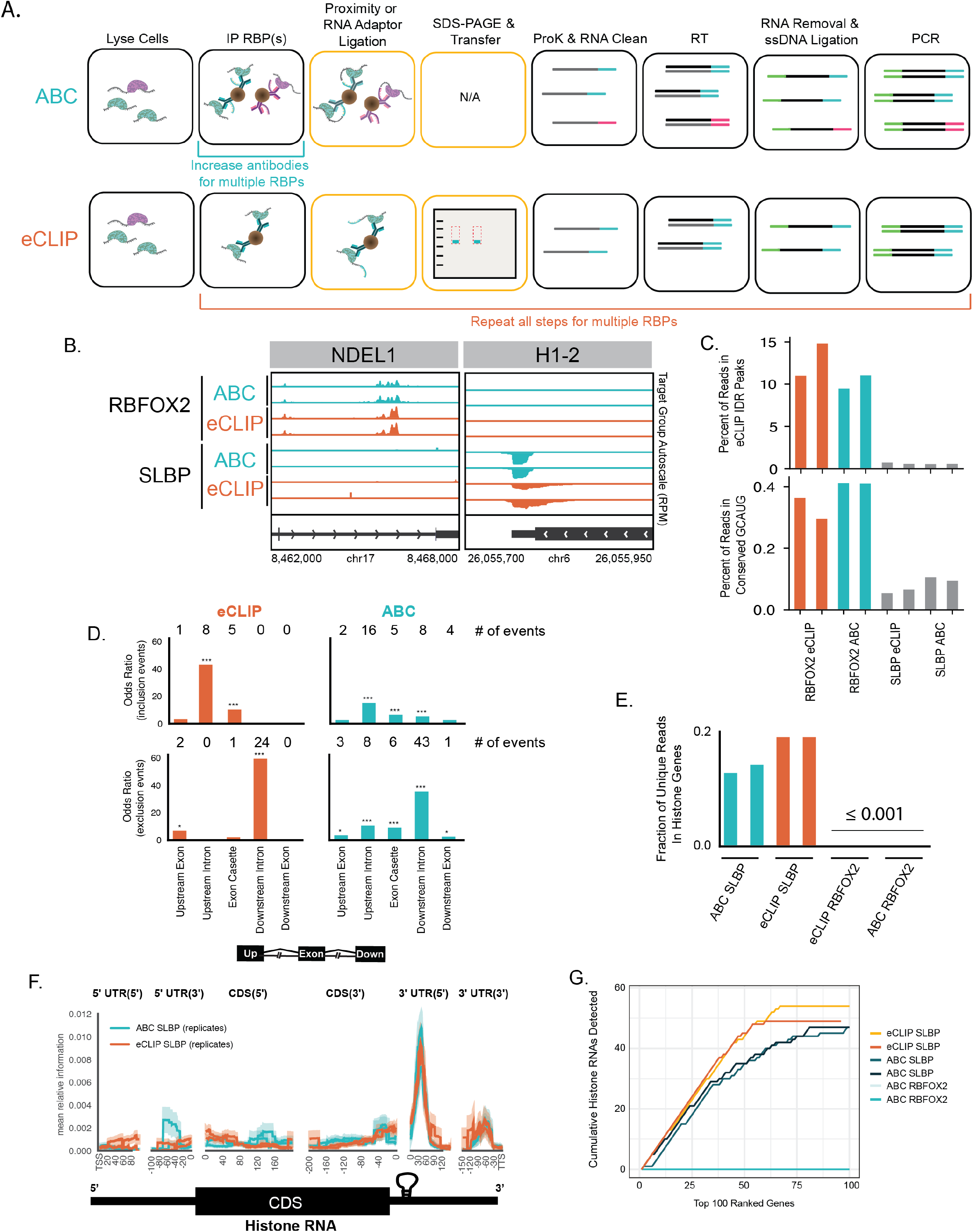
a) Schematic of ABC and eCLIP workflow. Yellow blocks highlight the difference between the two protocols. Barcoded oligos (30 nucleotides) are conjugated to IP-grade antibodies using click-chemistry prior to immunoprecipitation. The oligo then acts as the 3’ adapter and undergoes proximity-based ligation to RNA targets bound to the IPed RBP allowing for bioinformatic separation post-sequencing. b) Genome browser tracks showing binding sites of RBFOX2 and SLBP. Each panel is group normalized by RPM value. c) Percentage of uniquely mapped reads that are within eCLIP IDR peaks (top). Percent of reads aligning to conserved GCAUG sites (bottom). d) Peak enrichment (-log_10_*P*>3, lg_2_FC>3) in RBFOX2-dependent skipped exon events, defined as exons alternatively included/excluded upon RBFOX2 shRNA KD (Nature Methods 2016) (* indicate *P*< 0.05, ** *P*<10e-3; \*\*\**P*<10e-4). e) Fraction of uniquely mapped reads that map to histone mRNA. f) Metadensity profile of reads from ABC or eCLIP that map to histone mRNA. g) Cumulative count of histone genes across the top 100 ranked genes based on enrichment. Highly abundant background RNAs (mitochondrial and snoRNAs) were filtered out of all datasets.

To evaluate ABC with a transcriptome-wide view, we initially focused on RBFOX2 and observed that peaks from ABC showed similar enrichments to downsampled eCLIP data in proximal and distal introns (Supplemental Fig.2a) and were significantly enriched for the RBFOX2 motif (Supplemental Fig.2b). Reproducible peaks obtained from irreproducible discovery rate (IDR) analysis of RBFOX2 eCLIP data serve as empirically defined, highly ranked RBFOX2 sites. We observed that the proportion of ABC reads present within reproducible RBFOX2 also mirrors eCLIP (**Fig.1c**), a measure of the specificity of the ABC method. We also compared the fraction of reads that contained the conserved GCAUG sequence, as evolutionarily sequence conserved RBFOX2 motifs are more likely to be authentic sites^25^ (Supplemental Note1). We observed that the fraction of reads that contain the conserved motif is similar (~0.38% for eCLIP, and ~0.4% for ABC; **Fig.1c**). As RBFOX2 exhibits positional dependencies in its regulation of alternative splicing^25^, we demonstrated that ABC-derived peaks reproduced the eCLIP enrichment for RBFOX2 binding upstream and within exons that are included in the mature mRNA exons; as well as binding downstream to enhance exon recognition and exclusion from mature mRNA (**Fig.1d**). Next, we shifted our focus to SLBP. Both ABC and eCLIP displayed a similar fraction of reads that map to histone RNAs (**Fig.1e**). Metagene analysis also revealed a sharp peak at the well-characterized stem-loop within the 3’UTR of histone mRNAs (**Fig.1f**). To compare the gene level enrichment of both ABC and eCLIP, we ranked genes by the most enriched peaks after normalization and identified the top 100 genes in each dataset. Both technologies exhibited similar enrichment of histone genes (**Fig.1g**). Our comparison of ABC and eCLIP analyses for the RBPs RBFOX2 and SLBP suggests that ABC performs with comparable sensitivity and specificity to eCLIP at both read and peak-level features.

A defining advantage of ABC over current CLIP-based methodologies is that multiple RBPs can be interrogated simultaneously from a single sample (**Fig.2a**). To demonstrate this key functionality, in addition to RBFOX2, we selected nine other RBPs previously characterized by ENCODE III in K562 cells that exhibit a diversity of binding preferences within genic regions: DDX3 and EIF3G in the 5’ UTR; IGF2BP2, FAM120A, PUM2, and ZC3H11A in the 3’UTR; LIN28B in the CDS; SF3B4 involved in branch point recognition at the 3’ splice site; and PRPF8 which is downstream of the 5’ splice site. We performed replicate, multiplexed ABC experiments after conjugating barcoded oligonucleotides to each antibody raised against a specific RBP. These antibodies were previously validated and utilized in eCLIP analyses of these RBPs. After computational deconvolution of the barcodes, we processed each RBP within each ABC sample separately. For each RBP, we removed ABC reads that map to repetitive elements, only retaining reads that mapped uniquely to the human genome and performed peak-calling. We computationally downsampled the uniquely mapping eCLIP reads to the same sequencing depth as the ABC libraries (Supplemental Table1). We then performed peak-calling on the eCLIP samples. The numbers of initial peaks were similar between ABC and eCLIP (Supplemental Fig.3).

**Figure. 2:**
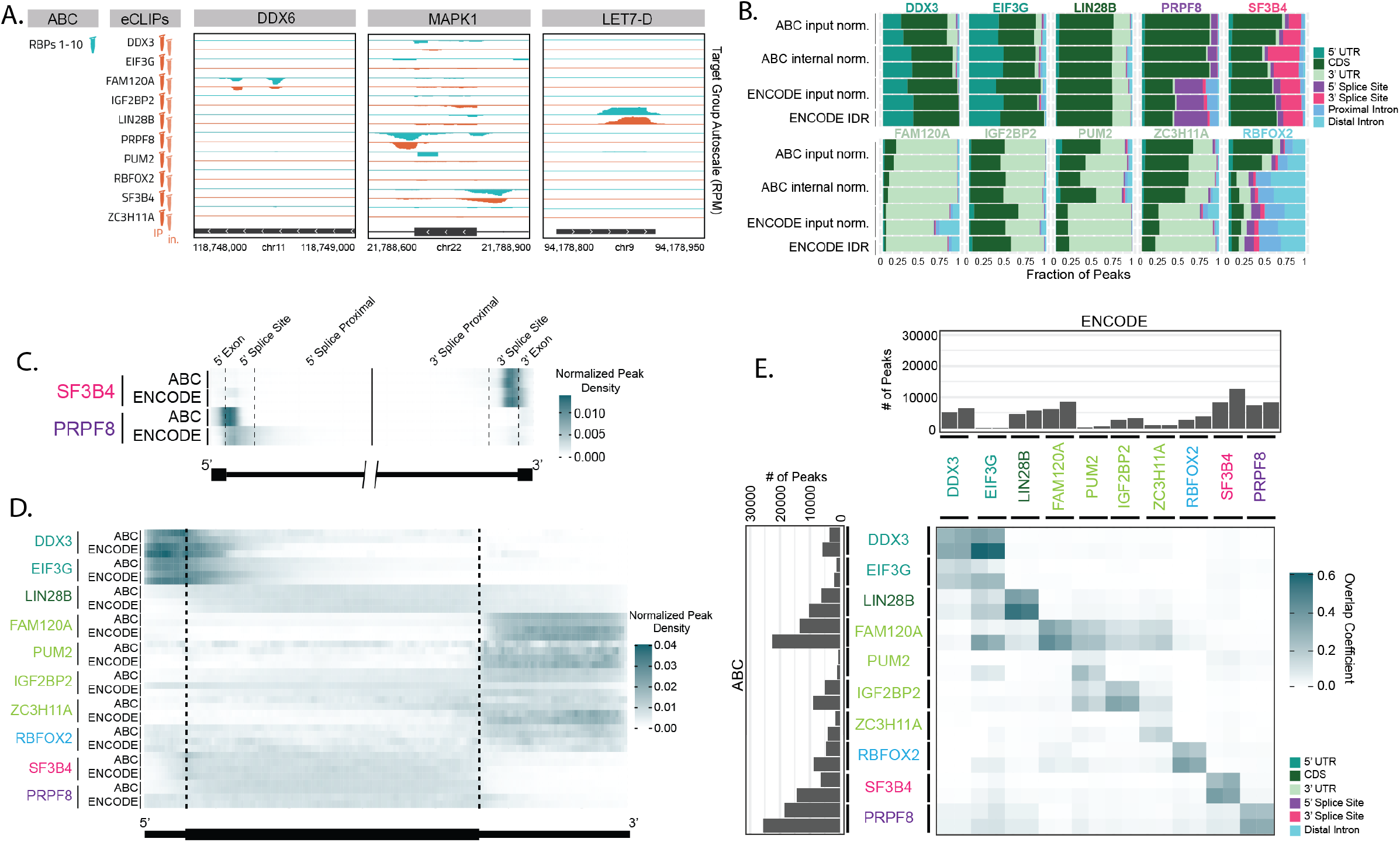
a) Genome browser tracks of select RBP binding sites depicting similar coverage between ABC (teal) and ENCODE eCLIP (orange). Each binding site was group normalized to all RBPs using RPM. b) Stacked bar plots of the fraction of peaks localized to each coding RNA feature in K562 cells. RBPs are color-coded with their annotated binding feature. ABC input normalization was compared to total RNA-seq. ABC internal normalization was a chi squared test between the other 9 RBPs. ENCODE input normalization was compared to its respective SMI. ENCODE IDR peaks were not downsampled. c) Splicing metagene profile of the two splicing factors SF3B4 and PRPF8 with density representing peak calls based on internal normalization (ABC) and input normalization (ENCODE). Peak intensity was normalized such that the total density for each sample was equal to 1. d) Internal normalized peaks (ABC) and input normalized peaks (ENCODE) were mapped across a normalized mRNA transcript for each RBP. Peak intensity was normalized such that the total density for each sample was equal to 1. e) Internal normalized peaks (ABC) and input normalized peaks (ENCODE) were intersected to find the number of overlapping peaks. The overlap coefficient is defined as (# overlapping peaks / total number of peaks in the smaller of the two datasets). The total number of peaks are displayed as a bar chart outside of the heatmap.

We then prioritized enriched peaks from ABC using total RNA-seq as background and compared them to ENCODE eCLIP datasets using the RBP’s size-matched input (SMI) control (Supplemental Note2). ABC produces comparable peak distributions across coding RNA features when compared to eCLIP (**Fig.2b**) as well as when analyzing all RNA features (Supplemental Fig.4). The eCLIP protocol incorporates a SMI to capture non-specific, background RNAs that are sequenced in a CLIP experiment. SMI allows for a measure of the experimental background in the CLIP experiment rather than providing an enrichment score relative to total RNA. As ABC removed the gel and membrane transfer steps, we reasoned that using the nine other RBPs in the multiplex may serve as an alternative approach to prioritize sites that are specific for each RBP. For a given binding site for a specific RBP, we computed the chi-square statistic from a 2×2 contingency table using the observed number of on-target reads in the given RBP peak and the total number of reads for that RBP versus the background of the number of off-target reads from the other nine RBPs and the total number of reads for those respective RBPs. Peaks with a *P* value of less than or equal to 0.001 were deemed statistically significantly enriched. Our enrichment strategy produced a similar peak profile to eCLIP analysis using SMI to prioritize binding sites (**Fig.2b**) with a similar number of total peaks (Supplemental Fig.3). Additionally, for the two RBPs, RBFOX2 and PUM2, both of which have well-characterized motifs, HOMER was able to *de novo* detect their respective motifs in the ABC samples (Supplemental Fig.5). Therefore, we conclude that a single ABC library (from 1 tube) generates similar overall results to 10 separate eCLIP experiments (from 20 tubes).

To further compare peak locations between ABC and eCLIP, we first plotted the metagene profiles of the enriched peaks for the spliceosomal proteins SF3B4 and PRPF8. Both RBPs displayed strong positional preferences proximal to their respective splice sites (**Fig.2c**). We observed that the ABC-derived peaks for PRPF8 were closer to the annotated 5’ splice sites than the eCLIP-derived peaks, resulting in changes to peak annotation **(Fig.2b**). All ten RBPs also displayed similar binding distributions in the metagene profiles (spliced mRNA) for both ABC and eCLIP (**Fig.2d**). Finally, to confirm that ABC and ENCODE were recovering the same binding sites, we computed the overlap coefficient between ABC and eCLIP replicates. There is a notable overlap between identified and enriched peaks in ABC and eCLIP (**Fig.2e**). In addition to intra-RBP reproducibility, there was overlap between RBPs known to bind similar features, like the 5’UTR binding proteins DDX3 and EIF3G. Average coverage of eCLIP peaks was also found to be correlated for all RBPs (Supplemental Fig.6). Finally, we wanted to evaluate if multiplexing RBPs had any appreciable effect on the quality of the data. No differences in peak distributions or quantity were observed when accounting for differences in read depth and peak coverage correlated between single and 10-plex ABC experiments (Supplemental Fig.7).

We conclude that ABC can effectively characterize the transcriptome-wide RBP binding sites for multiple RBPs from the same input amount as a single eCLIP experiment. This new protocol does not require an SDS-PAGE gel and generates data of comparable quality to eCLIP. Additionally, using a computational strategy to identify peaks that were enriched for specific RBPs within the pool, ABC obviated the separate SMI library requirement. These advantages result in at least a 20-fold reduction in the number of libraries generated for a single ABC 10-plex experiment. By simply increasing the number of barcodes, this advantage will grow. This unprecedented scalability will facilitate the broad annotation of RBPs in clinically relevant samples, like disease tissues, where source materials are rare and often input-limited.

## Acknowledgements

G.W.Y. is supported by NIH grants (HG004659, HG009889) and by an Allen Distinguished Investigator Award, a Paul G. Allen Frontiers Group advised grant of the Paul G. Allen Foundation. The authors would like to thank Julianne Dessert for creating the schematic artwork.

## Contributions

ABC was invented and initial proof-of-concept experiments and analysis were performed by D.A.L and K.B.C. Subsequent experiments were performed by D.A.L. and A.C.N. with bioinformatic analysis by D.A.L., K.A.S., H.H., K.R.H., S.C.B., and S.A.M. The manuscript was written by D.A.L., H.H., K.R., and G.W.Y with input from all authors.

## Methods

### Cell Culture

K562 and HEK293 cells were cultured in DMEM media supplemented with 10% FBS following the standard tissue culture technique. Cell pellets were generated by washing 10 cm plates (~15 million cells) once with cold 1X phosphate-buffered saline (PBS) and overlaid or resuspended with minimal (3 mL per 10 cm plate) cold 1X PBS and UV cross-linked (254 nm, 400 mJ/cm^2^) on ice. After cross-linking, cells were scraped and spun down, the supernatant removed, and washed with cold 1X PBS. Cell pellets (10 million each) were flash-frozen on dry ice and stored at −80°C.

### Antibody barcoding

100 µl of 100 µM oligo barcode (IDT) in PBS and 10 µl of 10 mM azide-NHS (Click Chemistry Tools cat# 1251-5) in DMSO were mixed at room temperature for two hours. Unreacted azide was removed by buffer exchanging into PBS using Zeba desalting columns (Thermo cat# 89883) following the manufacturer’s protocol. Azide labeled barcodes were stored at −20°C.

20 µg of antibodies were diluted to 70 µL in PBS, and the buffer was exchanged into PBS using Zeba desalting columns. 10 µL of 10 mM DBCO-NHS (Click Chemistry Tools cat# A134-10) was then added to the antibodies and allowed to rotate at room temperature for one hour ^26^. Unreacted DBCO-NHS was removed by buffer exchanging into PBS using Zeba desalting columns and stored at 4°C.

6.65 µL azide containing barcodes were then reacted with all the DBCO labeled antibodies (~70 µL). The mixture was allowed to rotate overnight at room temperature. Labeled antibodies were stored at 4°C and assumed to be 20 µg and used as is.

### Antibodies used in this study

| Antibody | Company | Catalog# |
| --- | --- | --- |
| RBFox2 | Bethyl | A300-864A |
| PUM2 | Bethyl | A300-202A |
| DDX3 | Bethyl | A300-474A |
| FAM120A | Bethyl | A300-899A |
| ACH11A | Bethyl | A300-524A |
| LIN28B | Bethyl | A300-588A |
| SF3B4 | Bethyl | A300-950A |
| EIF3G | Bethyl | A300-755A |
| PRPF8 | Bethyl | A300-921A |
| IGF2BP2 | MBL | RN008P |
| SLBP | Bethyl | A300-968A |

### Oligos used in this study

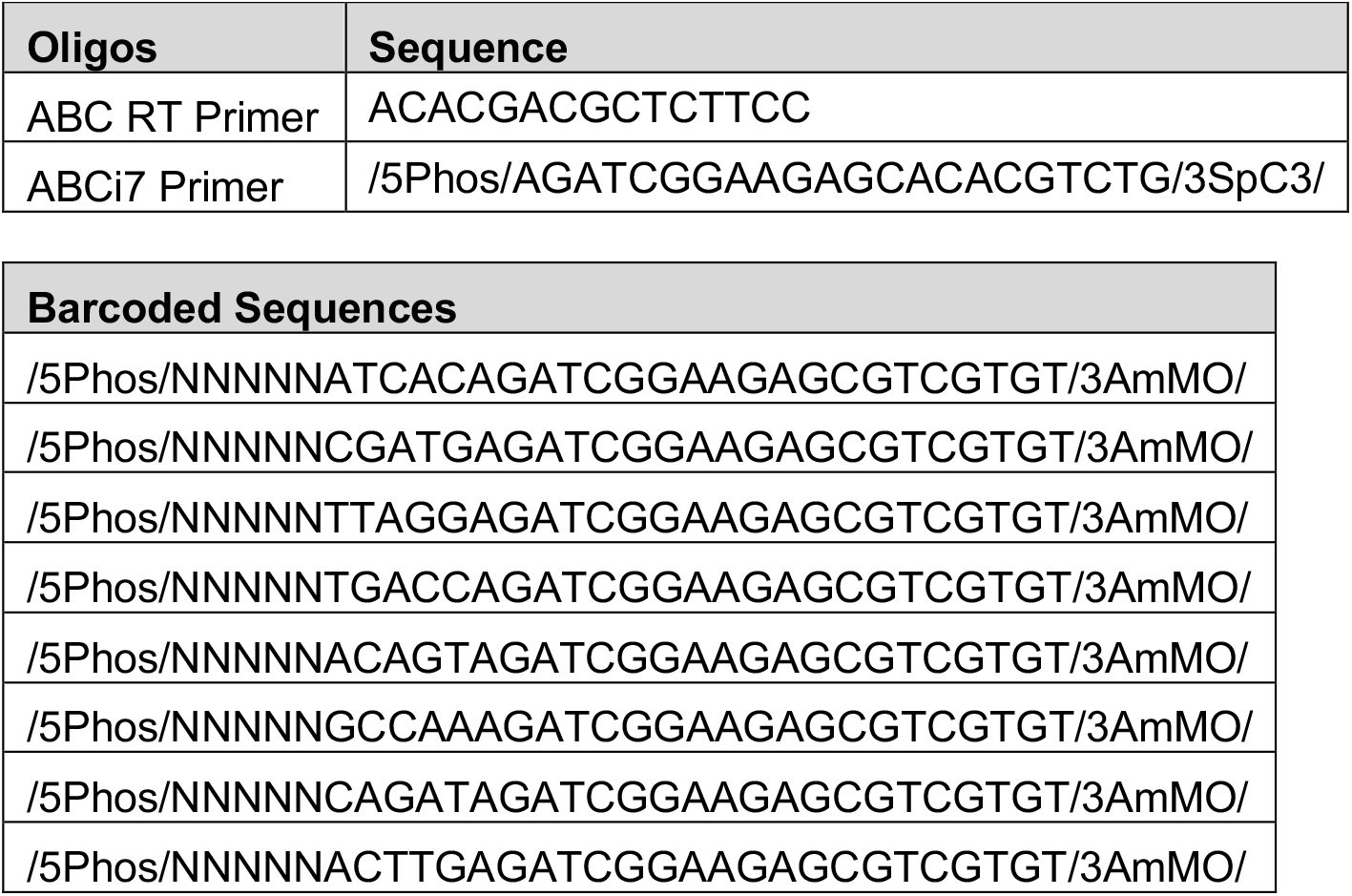
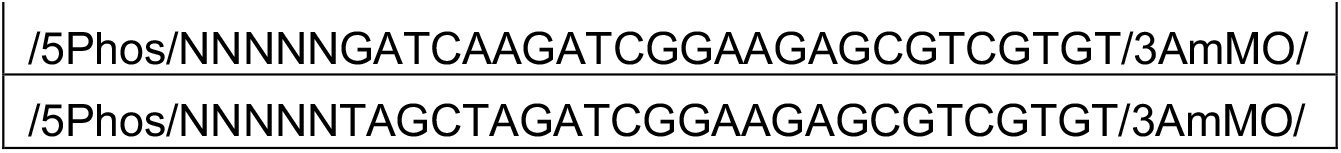

### Antibody conjugation CLIP

#### IP bead conjugation

200 µL of lysis buffer (50mM Tris pH 7.4, 100mM NaCl, 1% Igepal, 0.1% SDS, 0.5% Sodium Deoxycholate) was added to 25 µL anti-rabbit Dynabeads (Thermo Fisher cat# 11204D). Beads were washed twice with 500 µL lysis buffer before being resuspended in 50 µL lysis buffer. 5 µg of antibody was added and rotated at room temperature for one hour. Beads were again washed twice with 500 µL lysis buffer and resuspended in 50 µL lysis buffer. Repeat for each barcode and antibody combination.

#### Immunoprecipitation

Frozen HEK293 or K562 cell pellets were lysed in 1 mL lysis buffer supplemented with 5 µl protease inhibitor cocktail (Thermo Fisher cat# 87786) and 10 µl Rnase inhibitor (NEB cat# M0314B) and sonicated for 5 min with 30 second on/off cycles. The lysate was then treated with 10 µl diluted (1:25) Rnase I (Thermo Fisher cat# AM2295) and 5 µl TurboDNase (Thermo Fisher cat# AM2239B001) and incubated at 37°C for 5 min. Cellular debris was removed by centrifugation at 12,000 x g for 3 min. The supernatant was then transferred to a new tube along with 50 µL of each preconjugated antibody for each barcode (50 µL each from 10 different barcoded antibodies, 500 µL total for 10plex) coated magnetic beads and immunoprecipitated overnight at 4°C. Beads were subsequently washed with 500 µl high salt wash buffer (50mM Tris pH 7.4, 1M NaCl, 500 mM EDTA, 0.5% Igepal, 1% SDS, 0.5% sodium deoxycholate) (3x), high salt wash buffer + 80 mM LiCl (1x), and low salt wash buffer (500 mM Tris pH 7.4, 250 mM MgCl_2_, 5% Tween 20, 125 mM NaCl) (3x).

#### Proximity Ligation

Beads were resuspended in 80 µl T4 PNK reaction mix (3 µl T4 PNK (NEB cat# M0201B), 97.2 mM Tris pH 7, 13.9 mM MgCl_2_, 1mM ATP) and incubated at 37°C for 20 min with interval mixing. After PNK treatment, samples were washed with 500 µl high salt buffer (1x) followed by low salt buffer (3x). Proximity barcode ligations were carried out in 150 µl T4 ligation reaction mix (11 µL T4 ligase (NEB cat# M0437B-BM), 75 mM Tris pH 7.5, 16.7 mM MgCl2, 5% DMSO, 0.00067% Tween 20, 1.67 mM ATP, 25.7% PEG8000) at room temperature for 45 min with interval mixing. Samples were again washed with high salt buffer (1x) and low salt buffer (2x). Chimeric RNA barcode molecules were eluted from the bead by incubating with 127 µL ProK digestion solution (11 µL ProK (NEB cat # P8107B),100 mM Tris pH 7.5, 50 mM NaCl, 10 mM EDTA, 0.2% SDS) at 37°C for 20 min followed by 50°C for 20 min with interval mixing. Samples were placed on the Dynamagnet and supernatants were transferred to a clean tube. Samples were cleaned up with Zymogen RNA clean and concentrator following manufacturers protocol and eluted in 10 µL.

#### RT& Library Prep

RNA was reverse transcribed for 20 min at 54°C with 1.5 µL RT primer and 10 µl RT mix (0.6 µl Superscript III (Thermo Fisher cat# EP1756B012), 2.17x SuperScript III RT buffer, 10 mM DTT). After RT, excess primers and nucleotides were removed with 2.5 µl ExoSAP-IT (Thermo Fisher 75001.10.ML) and RNA was degraded with the addition of 1 µl 0.5 M EDTA and 3 µl 1M NaOH and heated at 70°C for 10 min, and pH neutralized with 3 µl 1M HCl. Samples were cleaned up using 5 µl MyOne Silane beads and eluted in 2.5 µl ssDNA ligation adapter (50 µL 100 µM ABCi7primer, 60 µL DMSO, 140 µL Bead Elution Buffer). Without removing the beads, 6.5 µl T4 ligase solution (76.9 mM Tris pH 7.5, 15.4 mM MgCl2, 3% DMSO, 30.8 mM DTT, 0.06% Tween 20, 1.5 mM ATP, 27.7% PEG8000), 1 µL T4 ligase, and 0.3 µL deadenylase (NEB cat# M0331B) was added and rotated overnight at room temperature. 45 µl bead binding buffer (0.001% Tween 20, 10 mM Tris pH 7.5, 0.1 mM EDTA) and 45 µl ethanol were added to the ligation mixture to rebind the cDNA to the silane beads. After washing with 80% ethanol the cDNA was eluted in 25 µl bead elution buffer and quantified by qPCR. Final libraries were amplified with dual index Illumina primers and sequenced on an Illumina Nextseq 2000.

### Data preprocessing

Data was processed similarly to the standard eCLIP pipeline^14^, except for a few adjustments to ABC’s multiplex design and library structure. For ABC data, UMIs were extracted using umitools 1.0.0^27^, and adaptors were removed using cutadapt 2.8^28^. Fastqs files were demultiplexed based on the 5’ nucleotide barcode sequence using fastx toolkit (http://hannonlab.cshl.edu/fastx_toolkit/). ABC libraries were sequenced on the reverse strand. Therefore, reads were reverse complemented before alignment to repetitive regions, removal of multi-mapped reads, and alignment to the genomic sequences using STAR 2.7.6. The pipeline is available at https://github.com/algaebrown/oligoCLIP.git.

### Calculating enrichment of peaks across different background inputs

After genome alignment of the ABC libraries, we used CLIPper (https://github.com/YeoLab/clipper) to call peaks using the immunoprecipitated library. Briefly, CLIPper uses nearby regions (+/-500 b.p.) to estimate the background in the immunoprecipitated library, followed by a Poisson distribution to assess the significance of a peak. We then ranked the enrichment of peaks across several different types of backgrounds, including: SMI (eCLIP), RNA-seq (ABC), or to other multiplex libraries (internal normalization). In the eCLIP protocol, the SMI library was prepared the same way as the IP library and captures the background cross-linking rate of other RBPs with similar size ranges as well as the RNA expression level and all other bias in library preparation. Since ABC does not have a SMI, we reasoned that a total RNA-seq library can capture part of the bias. Furthermore, internal prioritization to other RBPs within the multiplexed library can estimate not only the expression level, but also other biases introduced within the protocol. Ideally, as the number of RBP approaches infinity, the summation of “other RBPs in the multiplexed library” should approach the SMI-input. To estimate the significance of the peaks based on backgrounds, we used a chi-square test (or Fisher Exact test, if the number of reads < 5) to compare the number of reads in the IP library and the background library within and outside of the peak region. The pipeline is available at https://github.com/algaebrown/oligoCLIP.git.

### Evaluating the authenticity of peaks called

Since there exists no large-scale gold-standard standard datasets of binding sites, we made assumptions based on previous knowledge of certain RBPs. RBFOX2 has a strong binding to the GCAUG motif and its sites exhibit high sequence conservation across vertebrate evolution (which we operationally define as GCAUG sequences with phyloP > 3, in intronic and UTR regions) ^29^. RBFOX2 is also known to be enriched proximal to its regulated exons which exhibit positional dependent alternative splicing. We utilized the splicing microarray-defined (n=150) differentially included and skipped cassette exon events upon loss of RBFOX2^14^.

SLBP has been characterized to primarily bind stem loops within histone-encoding mRNAs. Based on these observations, we curated a set of “true positive regions” and assessed the sensitivity and specificity of our methods using AUROC and AUPRC. In addition, we compared our dataset of published eCLIP datasets in ENCODE. We calculated distribution of peaks in various transcriptomic features (UTR, CDS, intron, noncoding regions) and showed that the two methods capture a similar distribution for each RBP.

### Estimating library complexity

Library complexity (e.g. the number of unique molecules captured in the experiment) is a function of efficiency of every step within the library preparation as well as the sequencing depth. To ensure ABC has the same efficiency in capturing uniquely bound RNAs, we estimated at various sequencing depth of uniquely mapped reads, how many UMIs can be captured (Supplementary Fig.1a). Uniquely mapped reads were downsampled to various depths, then followed the preprocessing pipeline to deduplicate the reads.

### Exon Exclusion/Inclusion

The upstream exon is defined as the exon 5′ to the cassette exon. The up-flanking intron is defined as the intron between the upstream exon and the cassette exon. Whereas, the downstream exon is defined as the exon 3′ to the cassette, and the downstream flanking intron is the intron between the cassette and downstream exon. We then defined the background exons by randomly sampling 1500 exons with no change upon RBFOX2 KD. The odds ratio was calculated as: [(the number of skipped exons that overlapped with significant peak)/(the number of skipped exons that do not overlap with significant peak)]/[(the number of exons that overlap with background exons)/(the number of exons that do not overlap with background exons)]. “Overlap” is defined as at least 50% of peak length falling into the designated region. A Chi-square test is performed to test for significance in enrichment.

## Data Availability

All code for analysis is accessible here https://github.com/algaebrown/oligoCLIP.git. Data available at GEO accession: GSE205536.

## Supplemental Note 1. Calculation of the conserved motif

Conserved motifs within 3’UTR or intronic regions contained a score phyloP score > 3.

## Supplemental Note 2. Normalization of background and use of inputs in ABC/eCLIP comparison

We then focused on comparing RBFOX2 in both ABC and eCLIP peaks transcriptome-wide. As the number of peaks detected is a function of the number of reads, we downsampled to match the number of unique molecules for each library. First, we asked whether we need to control for transcriptomic background, as the SMI in eCLIP protocol, and if so, what is an appropriate background for ABC? Since ABC does not have an INPUT library, we utilized public HEK293T rRNA depleted RNA-seq to normalize the peaks. Motif analysis showed canonical GCAUG motifs in all regions in both the normalized and unnormalized peaks, albeit the normalized peaks have a cleaner motif with a lower *P* value. When ranking the peaks with −log10*P* value from either unnormalized peaks or normalized peaks, in CDS regions, the normalized peaks have AUPRC in classifying peaks containing the canonical GCAUG motif. We concluded that for high noise regions such as the CDS and UTR region, the peak calling procedure benefits from having INPUT background control to distinguish bona fide binding side from random highly expressed regions. Therefore, for subsequent analysis, all ABC peaks are normalized to cell-line matched RNA-seq.

## Extended Data Figure Legends

**Supplemental Fig.1:**
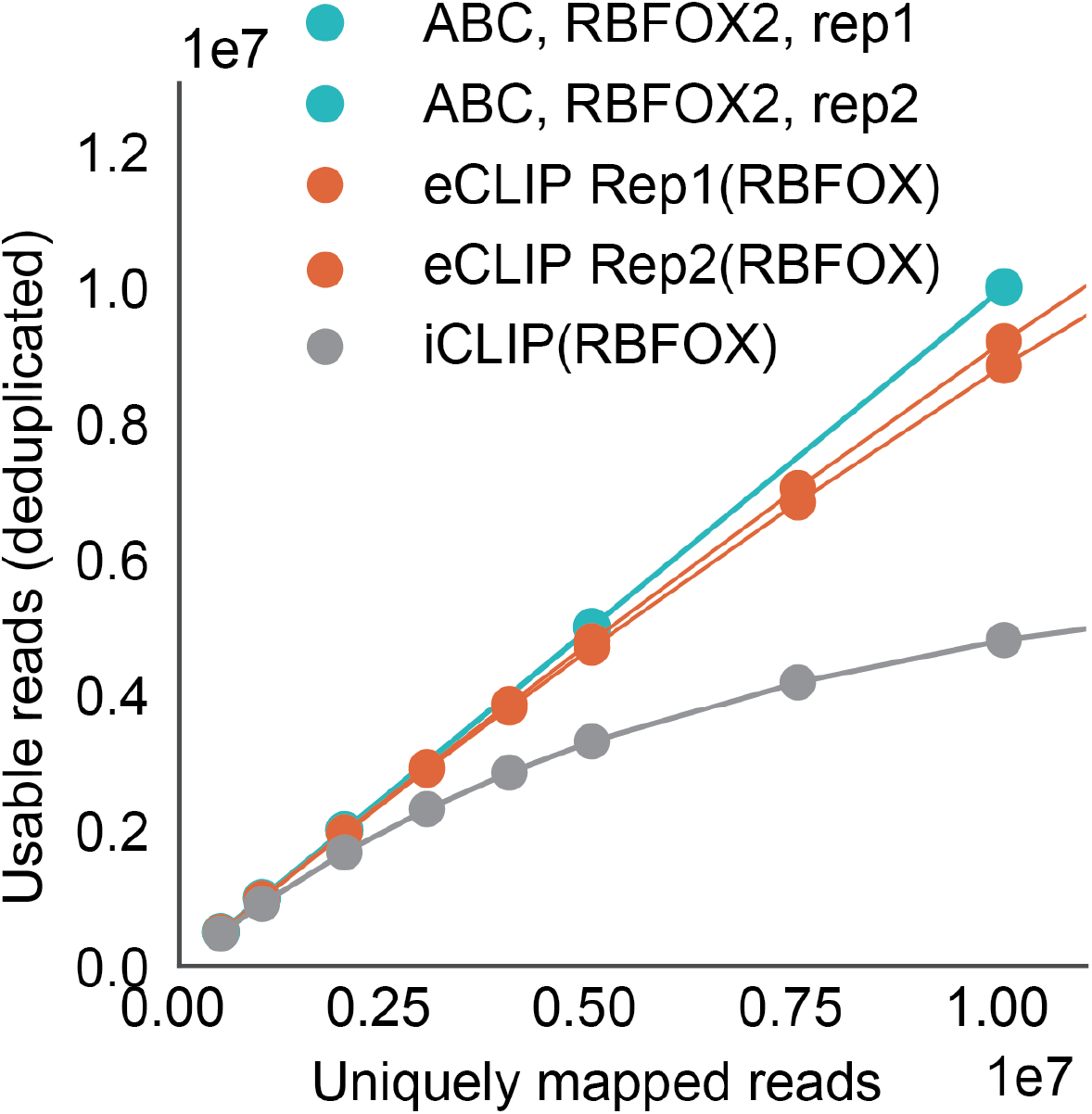
The number of unique molecules was estimated as a function of sequencing depth. After randomly sampling uniquely mapped reads from ABC, eCLIP, and iCLIP, we plot the number of uniquely mapped reads vs the number of usable reads.

**Supplemental Fig.2:**
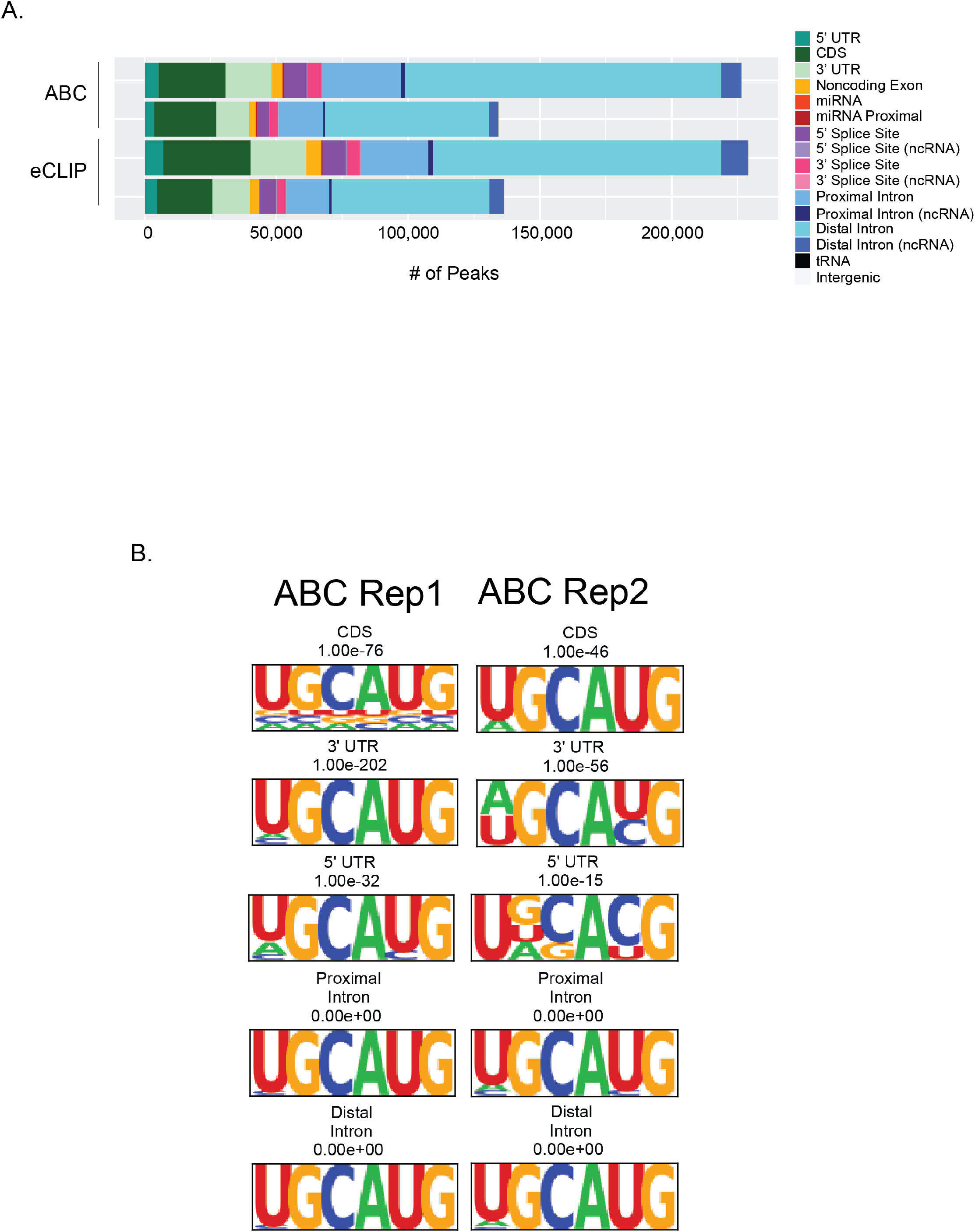
a) Stacked bar plots of the total number of peaks localized to each RNA feature from HEK293 cells. b) HOMER motif analysis of the significant peaks (-log10*P* value > 3, log2FC >3). Peaks were stratified by region (CDS, 3’UTR, proximal intron, or distal intron).

**Supplemental Fig.3:**
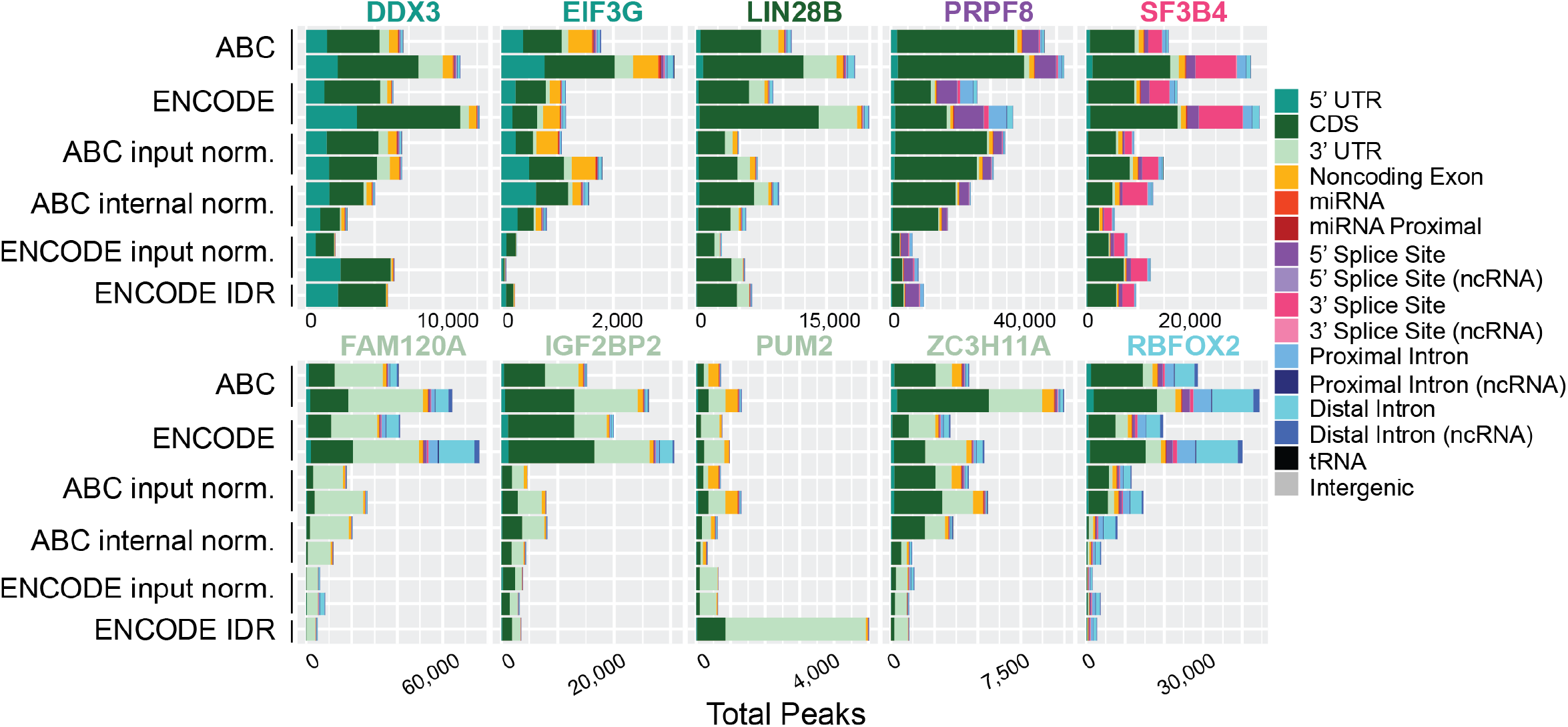
Stacked bar plots of the total number of peaks localized to each RNA feature in K562 cells. RBPs are color coded with their annotated binding feature. ABC input normalization was compared to total RNA seq. ABC internal normalization was a chi squared test between the other 9 RBPs. ENCODE input normalization was compared to its respective SMI. IDR peaks were not downsampled and used as is from ENCODE.

**Supplemental Fig.4:**
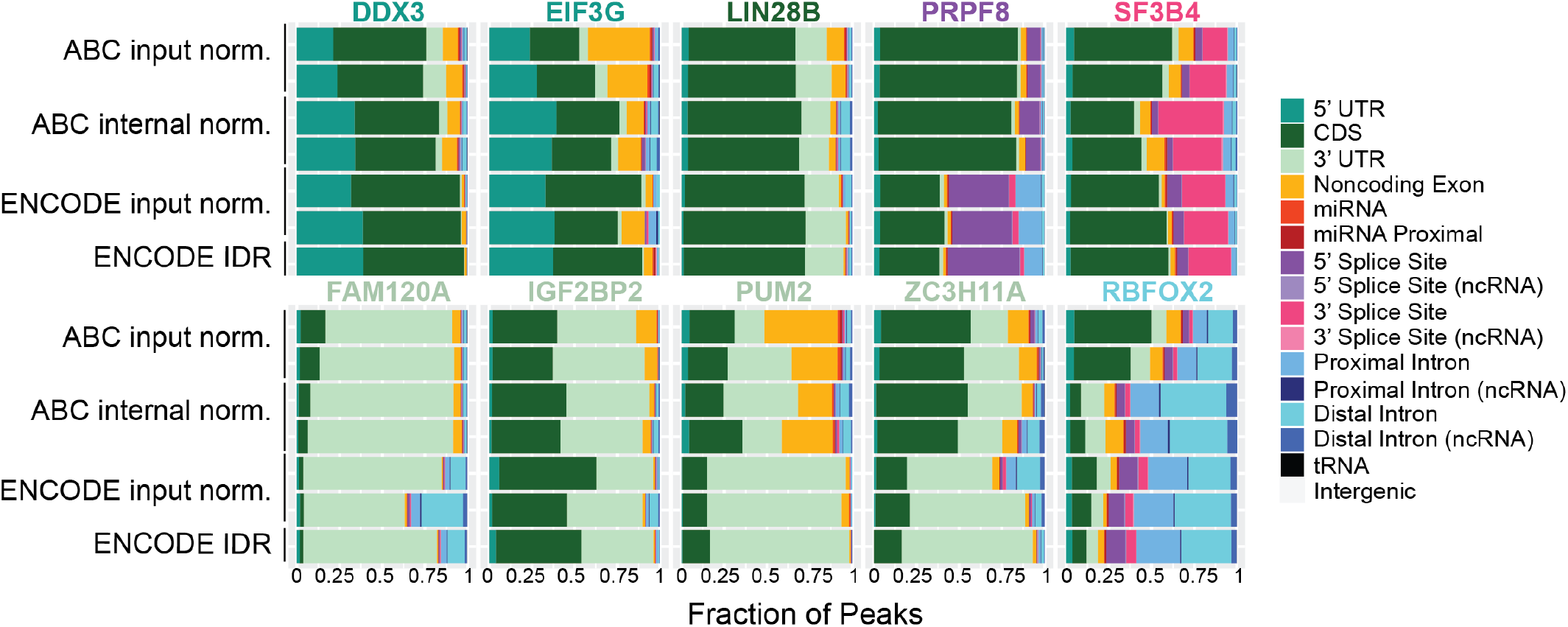
Stacked bar plots of the fraction of peaks localized to each RNA feature in K562 cells. RBPs are color coded with their annotated binding feature. ABC input normalization was compared to total RNA seq. ABC internal normalization was a chi squared test between the other 9 RBPs. ENCODE input normalization was compared to its respective SMI. IDR peaks were not downsampled and used as is from ENCODE.

**Supplemental Fig.5:**
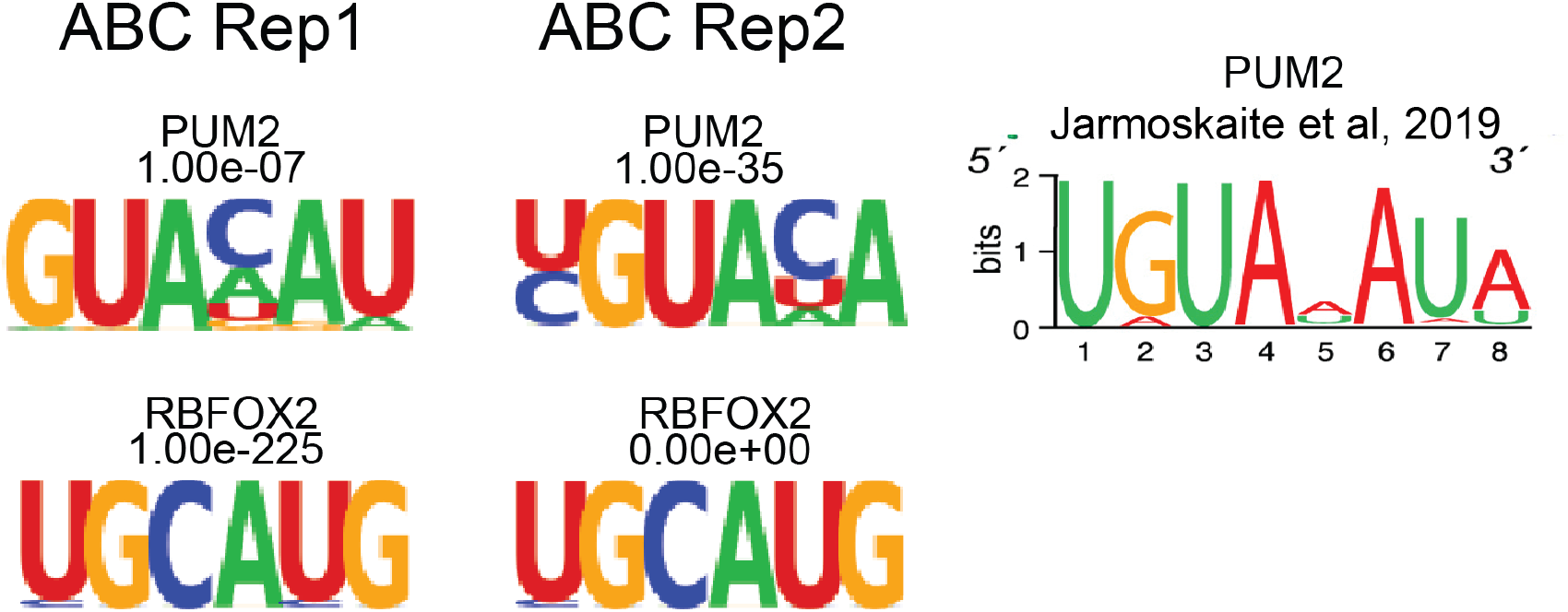
De novo motifs detected for PUM2 and RBFOX2 call from ABC internal normalized peaks. *P* values are listed for each RBP and sample. Reference PUM2 motif is provided from Jarmoskaite et al 2019.

**Supplemental Fig.6:**
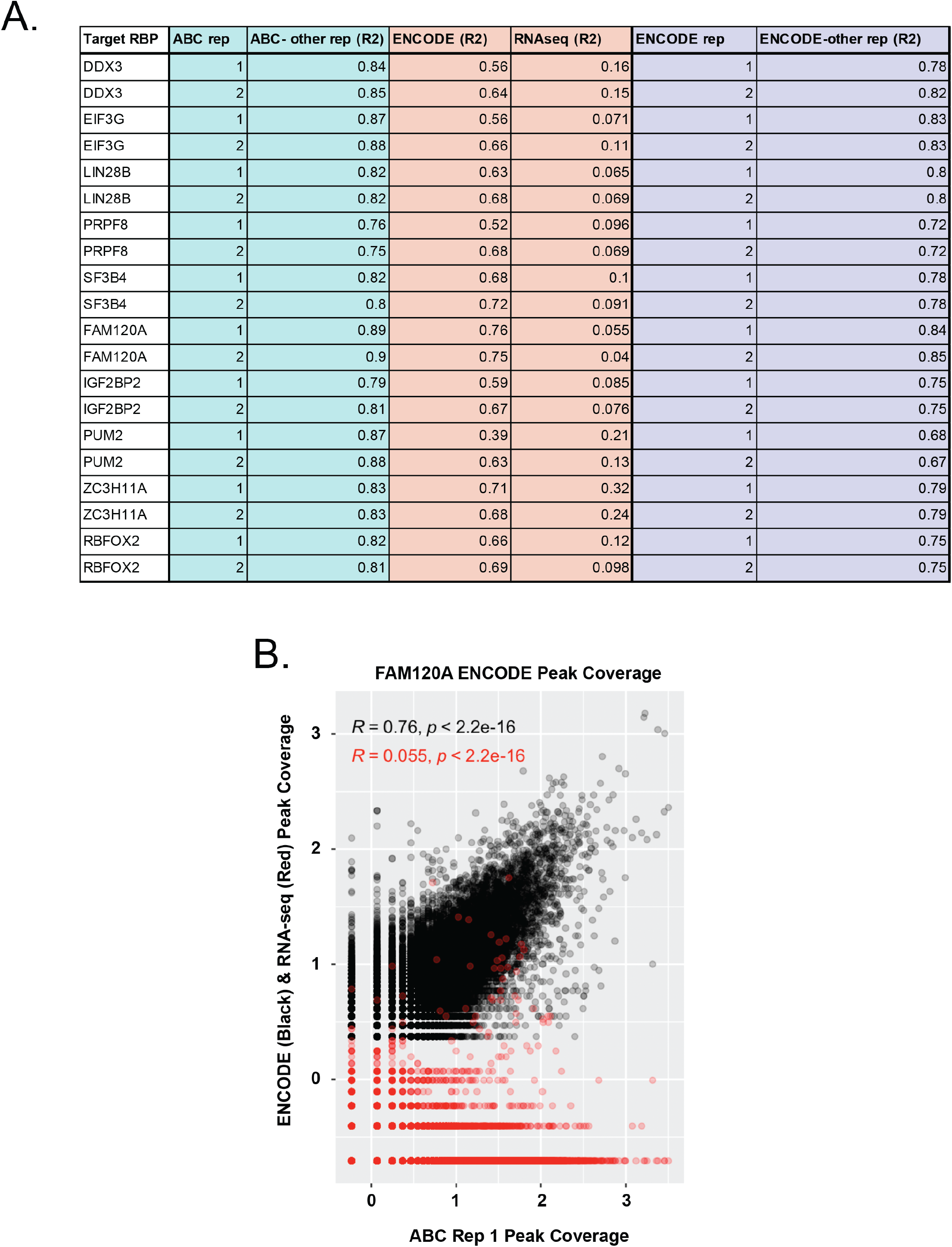
a) Table of Pearson correlation R^2^ values calculated from the coverage within ABC peaks (teal) and eCLIP peaks (orange and purple) between ABC, ENCODE, and RNA-eq experiments for each RBP. b) Example correlation plot of FAM120.

**Supplemental Fig.7:**
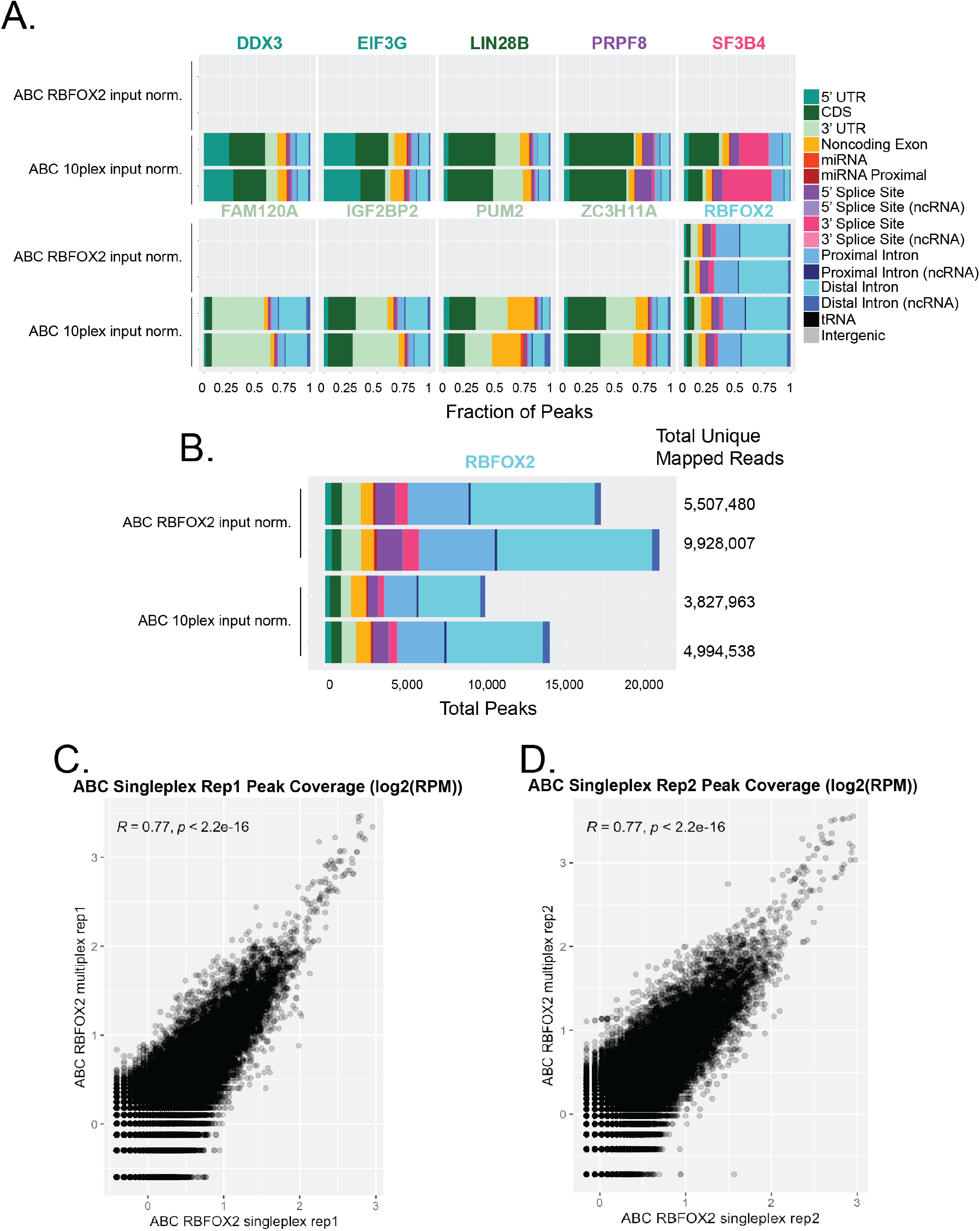
a) Stacked bar plots of the fraction of peaks localized to each RNA feature comparing single plex and multiplex ABC in HEK239 cells. b) Total number of peaks detected in single plex and multiplex ABC in HEK239 cells with total uniquely mapped reads listed on the right. c) Correlation of peak coverage between replicate 1 of simplex ABC vs multiplex ABC for RBFOX2. d) Correlation of peak coverage between replicate 2 of simplex ABC vs multiplex ABC for RBFOX2.

**Supplemental Table1:** Number of reads for each RBP in each 10-plex ABC experiment in K562 cells.

| **RBP** | **Rep** | **Repetitive Element**<br>**Reoved, Unique Genome**<br>**Aligned Deduplicated**<br>**Reads** |
| --- | --- | --- |
| DDX3 | rep1 | 460,353 |
| DDX3 | rep2 | 147,436 |
| EIF3G | rep1 | 146,496 |
| EIF3G | rep2 | 46,398 |
| FAM120A | rep1 | 1,636,905 |
| FAM120A | rep2 | 475,895 |
| IGF2BP2 | rep1 | 1,113,410 |
| IGF2BP2 | rep2 | 272,137 |
| LIN28B | rep1 | 347,579 |
| LIN28B | rep2 | 109,403 |
| PRPF8 | rep1 | 1,256,893 |
| PRPF8 | rep2 | 312,864 |
| PUM2 | rep1 | 67,591 |
| PUM2 | rep2 | 23,872 |
| RBFOX2 | rep1 | 916,888 |
| RBFOX2 | rep2 | 258,790 |
| SF3B4 | rep1 | 379,674 |
| SF3B4 | rep2 | 109,931 |
| ZC3H11A | rep1 | 205,252 |
| ZC3H11A | rep2 | 66,089 |

